# Condition-Dependent Noise Correlations without Condition-Dependent Spike Counts

**DOI:** 10.64898/2026.05.08.723078

**Authors:** Doyeon Kim, Matthew F. Panichello, Tirin Moore

## Abstract

The ability of the brain to encode information and control behavior depends on the coordinated activity of large and distributed neuronal populations. Correlations in neuronal spiking activity across trials of the same condition, or noise correlations (NCs), have been interpreted as a reflection of shared synaptic connectivity and as a contributing factor to the information capacity of neuronal populations. The impact of NCs on coding is most often considered in populations of neurons exhibiting robust condition-dependent information in their spike counts (SCs). However, theoretical work suggests that NCs could provide a source of condition-dependent information separate from SCs. We examined the activity of large neuronal populations in prefrontal cortex of macaques while they performed a spatial delayed response task composed of visual, memory, and motor epochs. We found that pairs of neurons that displayed visual, memory, and motor selectivity in their SCs often exhibited selectivity in their NCs, independent of spike count. However, we also found that pairs of neurons without SC selectivity during the different task epochs nonetheless exhibited condition-dependent NCs. Moreover, we found that the prevalence of condition-dependent NCs were largely comparable across neuronal pairs with or without SC selectivity. These results demonstrate that correlated variability in spiking activity can be condition-dependent even in the absence of condition-dependent SCs.

## Introduction

Representations of information by the brain have largely been characterized through the lens of neuronal firing rates. However, relying solely on trial-averaged firing rates offers an incomplete picture of population coding, as this approach assumes that neurons act as independent encoders, thereby ignoring the complex, moment-to-moment fluctuations that are ubiquitous in neural circuits (Panzeri et al., 2015). Information can be encoded not just in the mean rate but in the patterns of joint activity, which are captured by second-order statistics (Scaglione et al., 2011; Vaadia et al., 1995). For example, when an identical stimulus is presented repeatedly, trial-to-trial fluctuations in the response of one neuron are often correlated with those of others. These correlations represent shared variability that cannot be explained by the mean response to the stimulus (Ponce-Alvarez et al., 2013). Coordinated variability in the firing rates of populations of neurons has been linked to neural computations associated with myriad brain functions (Zavitz & Price, 2019), such as sensory coding (Bondy et al., 2018; Kohn & Smith, 2005; Singer, 1999), attention (Downer et al., 2017), and decision-making (Nassar et al., 2021). These correlations in neuronal spike counts (SCs), or noise correlations (NCs), have been interpreted as a reflection of shared synaptic connectivity and as a contributing factor to the information capacity of neuronal populations.

Previous studies have established that NCs among pairs of neocortical neurons often depend on the similarities in the selective properties of the constituent neurons (Hofer et al., 2011), consistent with the supposition that NCs reflect shared connectivity. In addition, other studies have revealed that for given neuron pairs, NCs also vary systematically across conditions, e.g. across different sensory stimuli (Ponce-Alvarez et al., 2013), an effect that could increase the information encoded in population activity that is complementary to neuronal firing rates (Josić et al., 2009; Pola et al., 2003; Panzeri et al., 1999; Shamir et al., 2004). However, to the extent that condition-dependent NCs contribute to population coding, that contribution has largely been considered in populations of neurons already conveying robust condition-dependent information in their firing rates, e.g. stimulus selective neurons (Franke et al., 2016; Zylberberg et al., 2016). Although condition-dependent NCs among populations of neurons without SC selectivity could also contribute to population coding, little or no empirical evidence of that dependence exists. However, the role of weakly- or non-SC selective neurons in neural computations and population coding, while less clear than that of SC selective neurons, has nonetheless become apparent in some neural systems. For example, in the mammalian visual system, the contribution of inhibitory interneurons to stimulus coding, e.g. orientation tuning, is well-established (Wood et al., 2018), yet these neurons are also known to be weakly tuned compared to excitatory neurons (Hofer et al., 2011; Kerlin et al., 2010; Runyan et al., 2013; Scholl et al., 2015), at least in part due to more extensive local connectivity (Hofer et al., 2011; Hu et al., 2012; Scholl et al., 2015). Similarly, neurons in prefrontal cortex exhibiting little or no SC selectivity during short-term memory tasks appear to contribute nonetheless to task performance (Aliramezani et al., 2025; Kim et al., 2016). Thus, it should be informative to assess the condition-dependence of NCs among neurons with non-selective SCs to various stimulus and task variables.

Building on these findings, we sought to determine the extent to which NCs provide a source of task-relevant information that is decoupled from individual neuronal tuning within the macaque lateral prefrontal cortex (LPFC). By recording from large neuronal populations during a spatial delayed response task, we investigated the degree to which NCs are modulated across distinct visual, mnemonic, and motor epochs. Specifically, we aimed to characterize the selectivity of NCs not only in neurons with robust SC selectivity but also in populations in which such selectivity is weak or absent. Below, we briefly summarize our results to date.

## Results

We analyzed the spiking activity of large populations of neurons from LPFC using Neuropixels probes of three macaque monkeys (Monkeys A, H, and J) (Panichello et al., 2024). During recordings, monkeys performed one of two versions of an oculomotor spatial delayed response task. In either version of the task, on each trial, one of eight possible peripheral locations was cued (Figure 1a), followed by a delay epoch during which the monkey needed to maintain the location in working memory. The trial concluded with a reward if the monkey made a saccadic eye movement response either to the remembered location or to one of two appearing targets that matched the cued location. In each recording session, we simultaneously measured the spiking activity of hundreds of neurons (mean ± SEM, 329 ± 46; total n = 8,225 units across 25 sessions) across LPFC (area 8/9/46, see Methods).

**Figure 1.**
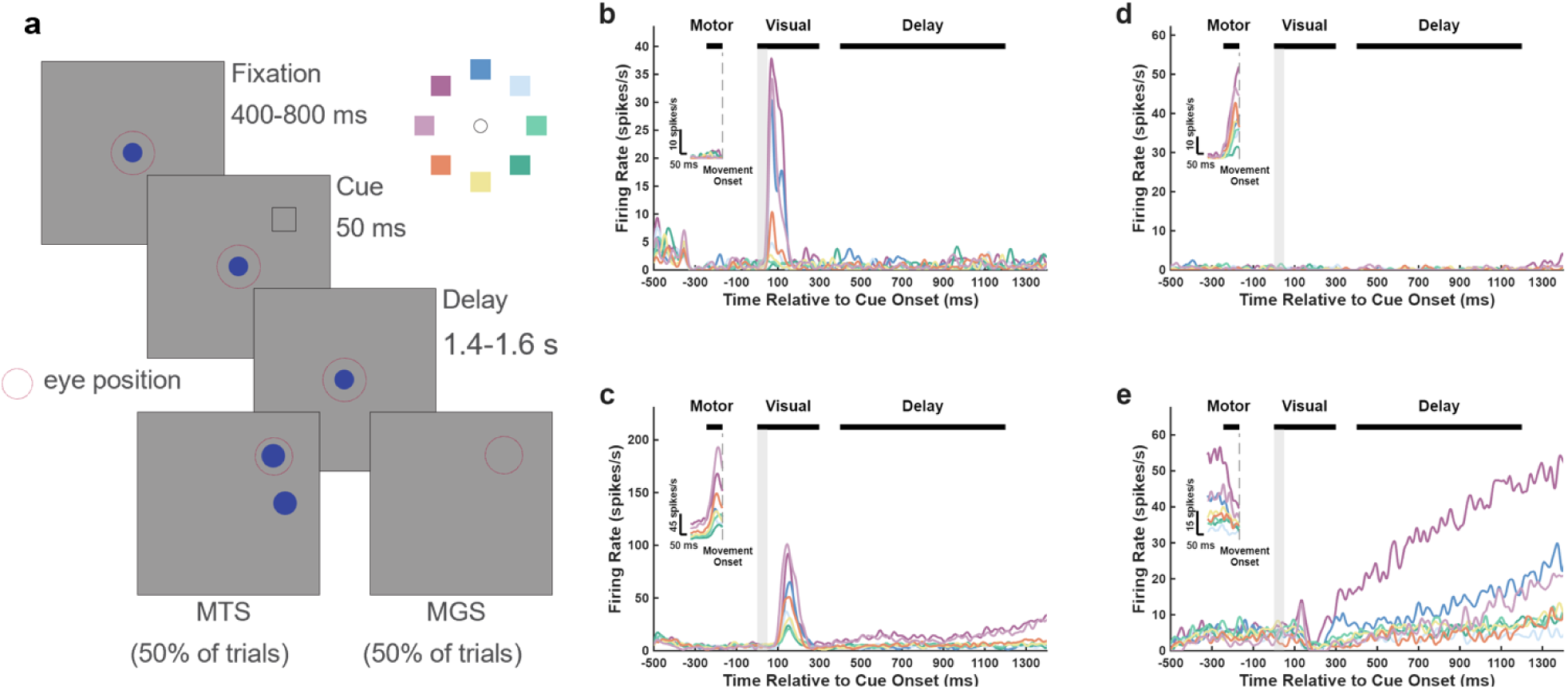
Task paradigm and functional diversity of prefrontal neurons. **a,** Schematic of the delayed Match-to-Sample (MTS) and Memory-Guided Saccade (MGS) tasks. On each trial, a cue was presented at one of eight possible locations (upper right inset). Following a memory delay epoch, the animal received juice as the reward for making an eye movement to the previously cued location (red circles indicate eye position). **b-e**, Trial-averaged peristimulus time histograms (PSTHs) for four example prefrontal neurons exhibiting diverse SC selectivity across task epochs: Visual, Delay, and Motor. Horizontal black bars at the top of each panel indicate the analysis windows for each epoch, and grey vertical bars indicate the cue presentation epoch (0-50 ms). Insets display the PSTHs aligned to movement onset during the Motor period (dashed vertical lines indicate movement onset). Example neurons with condition-dependent SC activity during the **b**, Visual epoch, **c**, Visual and Motor epochs, **d,** Motor epoch, **e**, Delay and Motor epochs. Cue locations are color-coded as shown in the inset of panel a.

We first assessed the SC selectivity of individual neurons in each of the task epochs by comparing their responses across cue locations during the presentation of the visual cue (“Visual”; 0-300 ms post-cue onset), memory delay (“Delay”; 400-1,200 ms post-cue), and the saccadic response (“Motor”; -125-0 ms relative to movement onset) (see Methods). As in previous studies (Bruce & Goldberg, 1985; Funahashi et al., 1989), individual neurons showed varying degrees of SC selectivity across the Visual, Delay and Motor epochs (Figure 1, b-e). Within each epoch, we identified SC selective neurons as those who drove successful classification of the cue location and whose ablation drove decoding performance down to chance-level (see Methods). A large proportion of neurons showed SC selectivity for each of the epochs (Visual, 47.424%; Delay, 44.286%; Motor, 57.005%). For example, the neuron shown in Figure 1b exhibited SC selectivity during the Visual and Motor epoch but was non-selective during the Delay epoch. Other neurons exhibited selective responses during different epochs, such as the Delay and Motor epoch (Figure 1e). These observations highlight the well-known observation that neurons do not necessarily display selectivity across all task epochs but can display different combinations of Visual, Delay, and Motor selectivity in their SCs.

Next, we measured the pairwise NCs (r_sc_) in simultaneously recording neurons across the full population. In many individual neuronal pairs, NCs were found to depend on the stimulus condition. For example, for the same pair of neurons, the NCs could be significantly positive in one condition and significantly negative in another. This was observed in each of the three trial epochs (Figure 2a). Next, to leverage the broad, and relatively continuous, range of distances between simultaneously recorded neuronal pairs, we examined the dependence of NCs on cortical distance (Figure 2b). Consistent with previous findings (Smith and Kohn, 2008), we found that |r_sc_| declined monotonically with increasing distance between neuronal pairs in all three epochs (Spearman’s rank correlation; Visual, r = -0.064, p < 0.001; Delay, r = -0.062, p < 0.001; Motor, r = -0.064, p < 0.001). The overall mean magnitude of |r_sc_| across all distances varied across epochs (Visual = 0.099; Delay = 0.131; Motor = 0.083) most likely reflecting the differences in the corresponding analysis windows, i.e. Motor<Visual<Delay (Cohen & Kohn, 2011; Smith & Kohn, 2008). Overall mean firing rates were similar across the three epochs.

**Figure 2.**
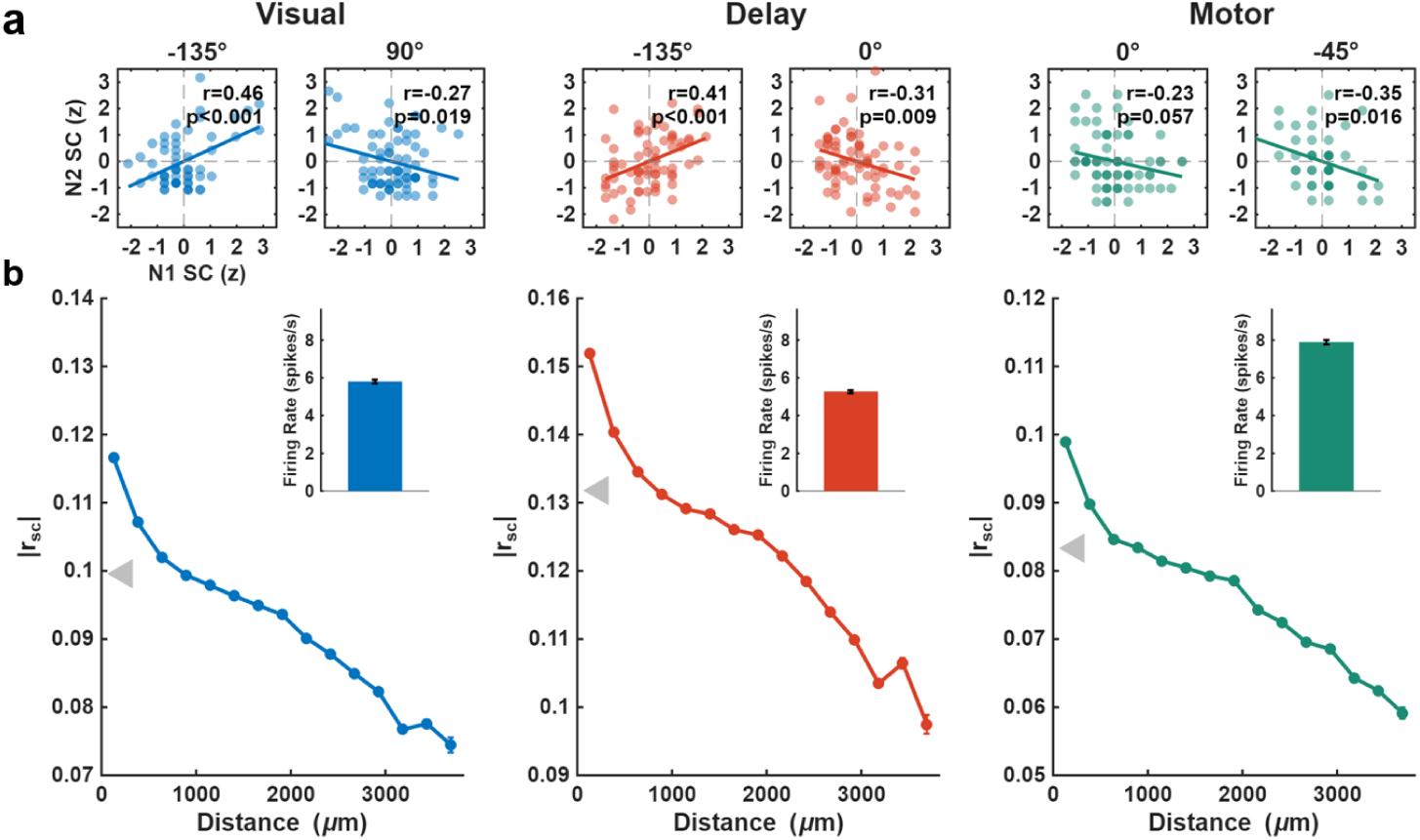
Pairwise NCs and their dependence on cortical distance. **a,** Examples of neuronal pairs exhibiting condition-dependent NCs. Scatter plots display trial-by-trial z-scored SCs for two neurons for different cue location conditions. Line-of-best fit, Pearson correlation coefficients (r), and p-values are indicated within each subplot. Angles indicate cue condition; 0° indicates the rightmost cue, values increase clockwise. **b,** Average absolute NC between neuronal pairs plotted against their cortical distance (μm) during the Visual (left), Delay (middle), and Motor (right) epochs. Error bars represent the standard error of the mean (SEM). The grey arrows indicate the overall mean of absolute NC. Insets in each panel show the mean firing rate (spikes/s ± SEM) of the corresponding neuron pool during each epoch.

We next assessed whether NCs were condition-dependent across the population by examining pairs of neurons with significant condition-dependent SC tuning (see Methods). Figure 3a illustrates a representative pair of SC selective neurons during the Delay epoch. Both neurons exhibited significant SC tuning across the eight different cue locations (p < 0.001). That is, their SCs were condition-dependent. However, the NCs between these neurons were not constant but varied systematically with the cue location, suggesting that NCs were also condition-dependent.

**Figure 3.**
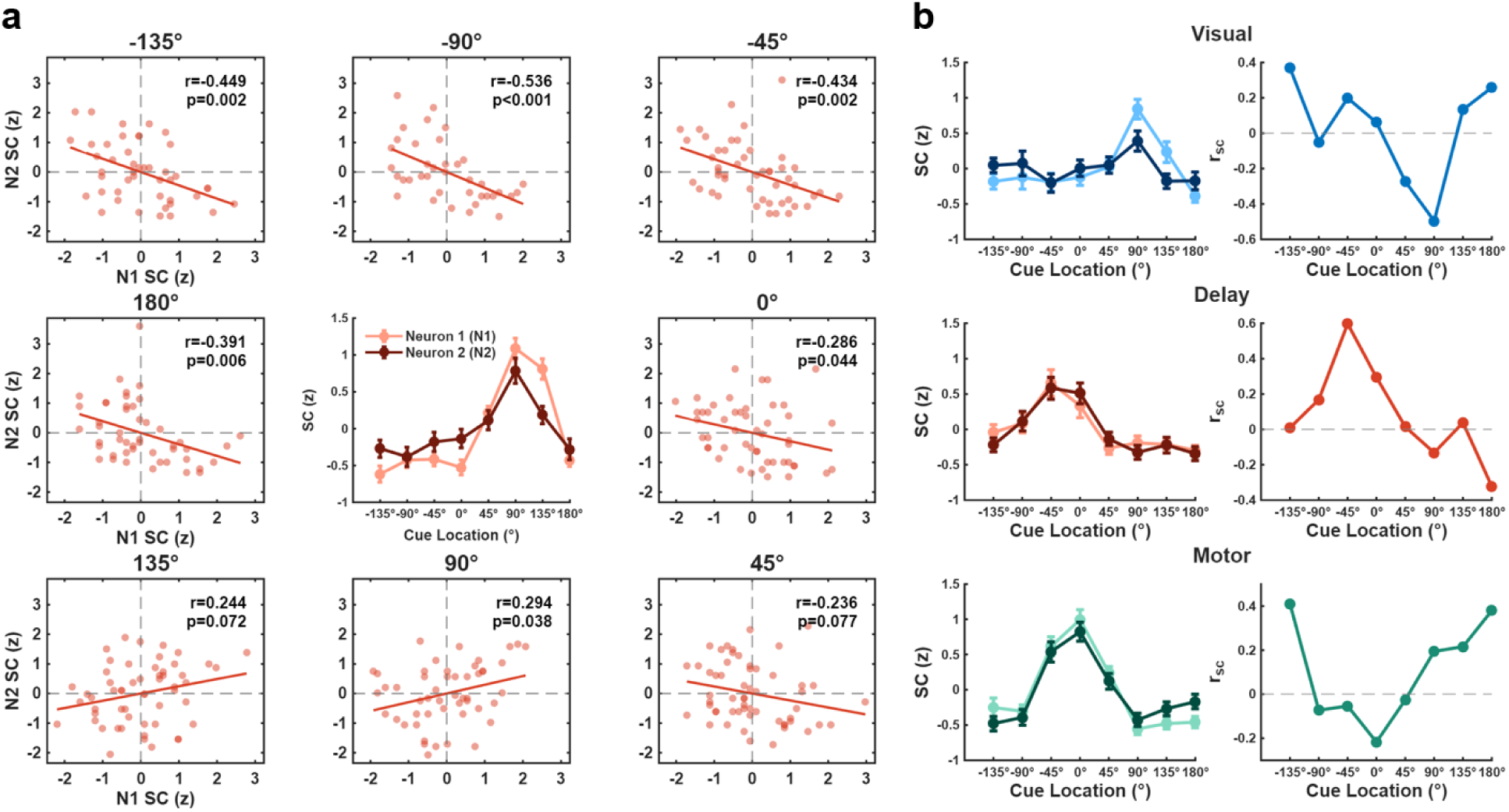
NCs and SC tuning curves for selective neuron pairs. **a,** Example of condition-dependent modulation of NCs in a selective neuron pair during the Delay epoch. The central subplot displays the SC tuning curves of two neurons across cue locations. Error bars represent standard error of the mean (SEM). Surrounding subplots show the trial-to-trial scatter of z-scored SCs for the two neurons at each corresponding cue location. Solid lines indicate linear regression fits. Pearson correlation coefficients (r) and p-values are indicated in the top right for each cue location. **b,** Examples of selectively tuned neuron pairs across the Visual (top), Delay (middle), and Motor (bottom) epochs. Left column: tuning curves with SCs (z-scored) of the two neurons across eight cue locations. Right column: Pairwise NC (r_sc_) values for each corresponding neuron pair from the left column, across the same cue locations.

To identify pairs in which NCs carried information about the cue, we z-scored the SCs for each neuron in a pair within each cue location to remove the effect of cue location on mean SC, then modeled their relationship using linear regression as:

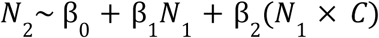

where N_2_ is the normalized SC of one neuron, N_1_ is the normalized SC of the other, and C is a categorical coding of cue location. A significant interaction term (p < 0.05) thus indicates that the NC between neurons is modulated by cue location. For example, for the neuron pair in Figure 3a, the regression analysis revealed a significant modulation of NCs by cue location during the Delay epoch (p < 0.001).

Next, we extended this analysis to different task epochs. As shown in the examples in Figure 3b, we observed pairs of neurons with condition-dependent SCs also exhibited condition-dependent NCs across the Visual, Delay, and Motor epochs. Our regression analysis revealed a significant modulation of NCs by cue location in each case (Figure 3b; Visual and Motor epochs, p < 0.001; Delay epoch, p < 0.05).

We next asked if condition-dependent NCs were confined to SC selective neurons by performing the above analyses on non-selective neuron pairs, defined as neurons whose removal did not further degrade decoding performance below chance-level (see Methods). Figure 4a shows two representative neurons that were non-selective during the Delay epoch. However, despite the absence of delay tuning, NCs in the delay period varied across cue location conditions. The interaction term in the regression model confirmed this condition-dependent modulation in NCs for this pair (p < 0.001). Other pairs displayed condition-dependent NCs in the Visual and Motor epochs (Figure 4b). For each example case in Figure 4b, regression analyses revealed a significant modulation of NCs by cue location (Visual, p = 0.043; Delay, p < 0.001; Motor, p =0.015).

**Figure 4.**
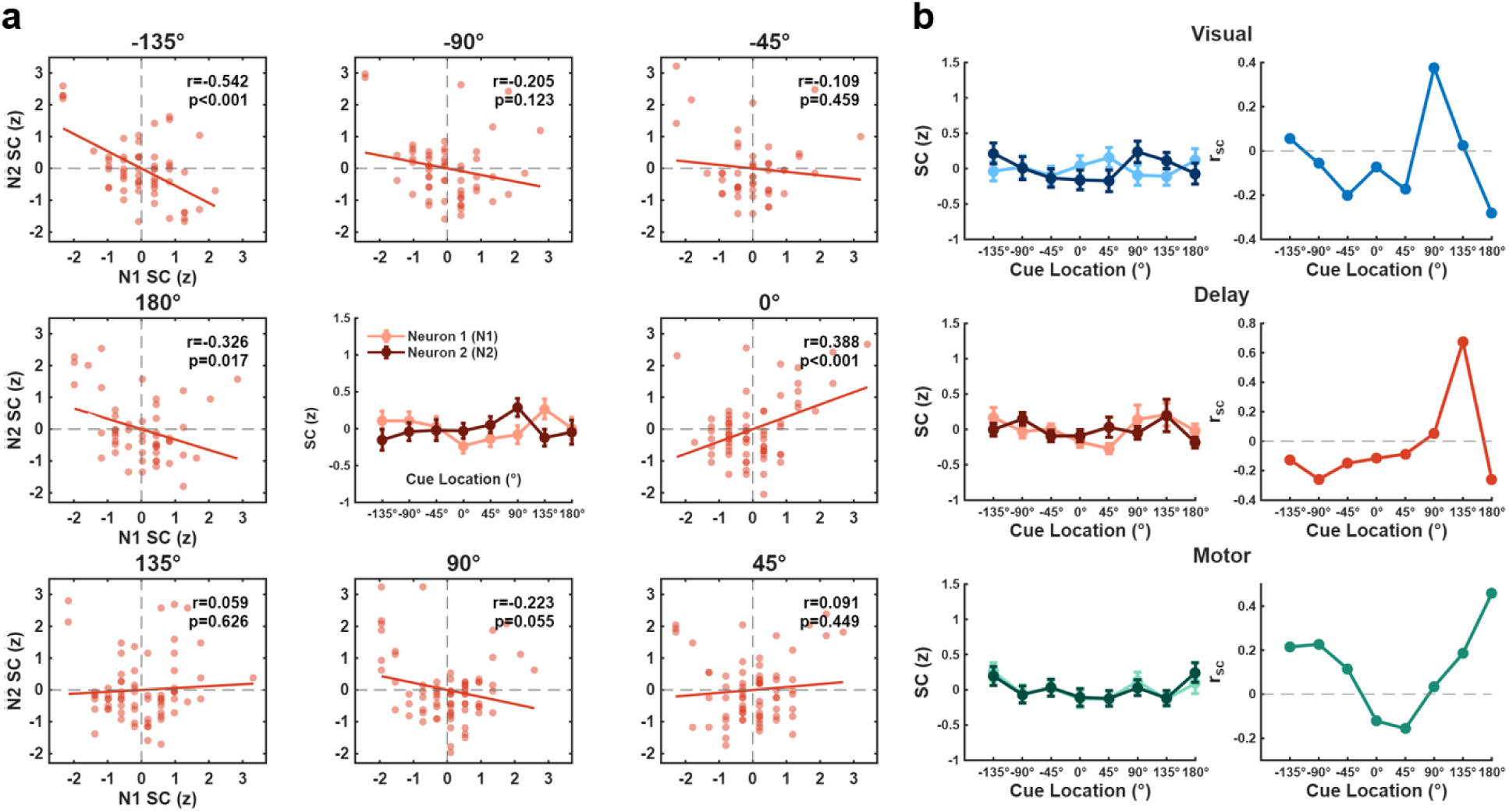
NCs and SC tuning curves for non-selective neuron pairs. **a,** Example of condition-dependent modulation of NCs in a non-selective neuron pair during the Delay epoch. The central subplot displays the SC tuning curves of two neurons across eight cue locations (error bars, SEM). Surrounding subplots show trial-to-trial scatterplots of z-scored SCs for each cue location. Solid lines indicate linear regression fits, with Pearson correlation coefficients (r) and p-values provided for each panel. **b,** Examples of non-selective neuron pairs across task epochs (top: Visual, middle: Delay, bottom: Motor). Left column: tuning curves with SCs (z-scored) across eight cue locations, showing relatively flat tuning for both neurons. Right column: Pairwise noise correlation (r_sc_) values for the corresponding pairs from the left column.

Having observed condition-dependent NCs in individual neuron pairs, we next assessed the extent to which that effect depended on the distance between neuron pairs. We hypothesized that since NCs typically scale with the distance between neurons (Shapcott et al., 2016; Shilling-Scrivo et al., 2022; Smith & Kohn, 2008) so too should the prevalence of condition-dependent NCs. Indeed, we observed a clear distance-dependent decline across all sessions (Figure 5). Linear regression analyses revealed that neurons in closer proximity were significantly more likely to share condition-dependent NCs across all three epochs (Visual, Delay, and Motor; all p < 0.001).

**Figure 5.**
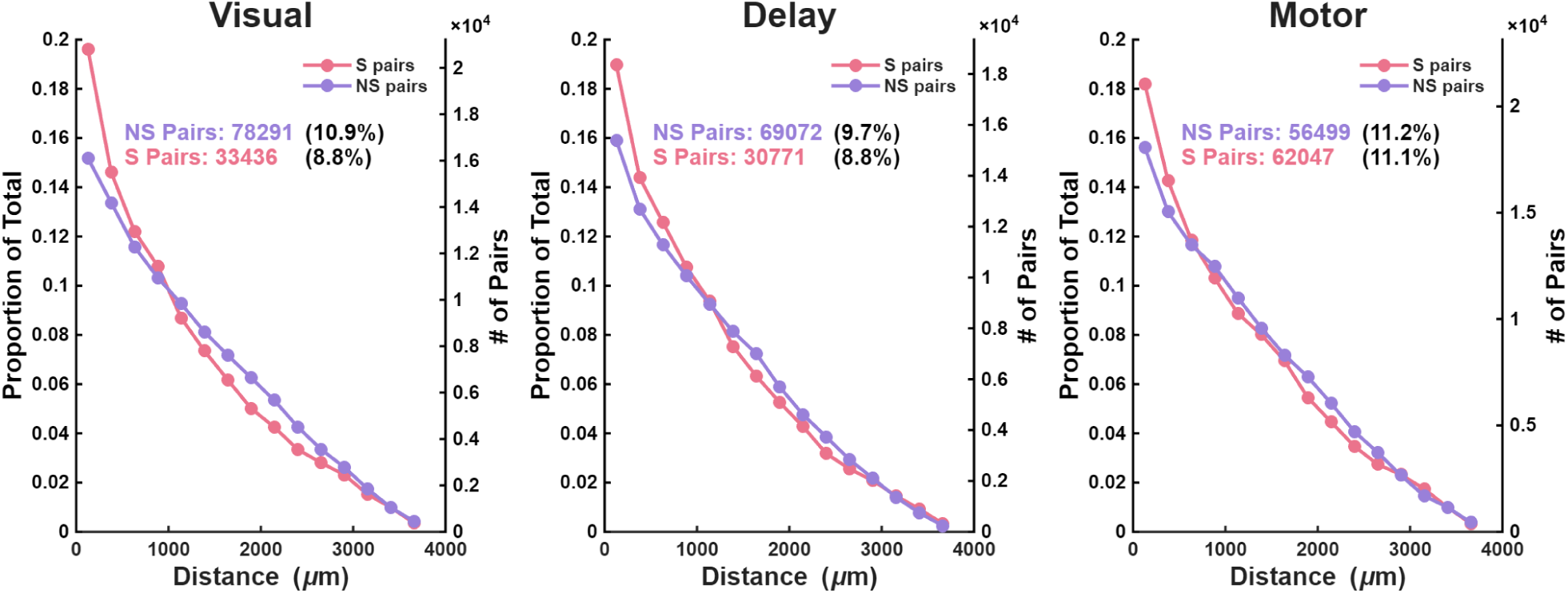
Condition-dependent NCs decay with increased distance between neuronal pairs. Prevalence of condition-dependent NCs across cortical distance for selective (S) pairs (pink) and non-selective (NS) pairs (purple) across the Visual (left), Delay (middle), and Motor (right) epochs. The left y-axis denotes the proportion of pairs with condition-dependent NCs, while the right y-axis displays the neuron pair counts. Text insets specify the total count and percentage of pairs with condition-dependent NCs.

Focusing on neuronal pairs separated by fewer than 1000 μm, we next quantified the prevalence of condition-dependent of NCs in neuronal pairs in which both neurons were selective, both were non-selective, or pairs in which one was selective and the other was non-selective (mixed). Specifically, we calculated the percentage of pairs with a significant condition-dependent interaction term in the regression model out of the total number of selective, non-selective, and mixed pairs (Figure 6, see Methods). Comparing across epochs, the proportion of condition-dependent NCs did not differ significantly between the Visual and Delay epochs for selective (p = 0.980) or mixed pairs (p = 0.992). In contrast, there was a greater percentage of condition-dependent NCs in the visual epoch for non-selective cells (p < 0.001). All 3 pair types showed the greatest proportion of condition-selective NCs in the motor epoch (all comparisons, p < 0.001) despite the smaller overall |r_sc_| values in that epoch (Figure 2b). Notably, when comparing across selective, non-selective and mixed neuronal pairs, we found that significant condition-dependent NCs were evident in all three types. During the Visual and Motor epochs, non-selective cells exhibited the highest proportion of condition-dependent pairwise NCs, whereas selective cells exhibited the highest proportion during the Motor epoch (Figure 6, all pairwise comparisons p < 0.001). These findings demonstrate that the prevalence of condition-dependent NCs was comparable across neuronal pairs with and without SC selectivity.

**Figure 6.**
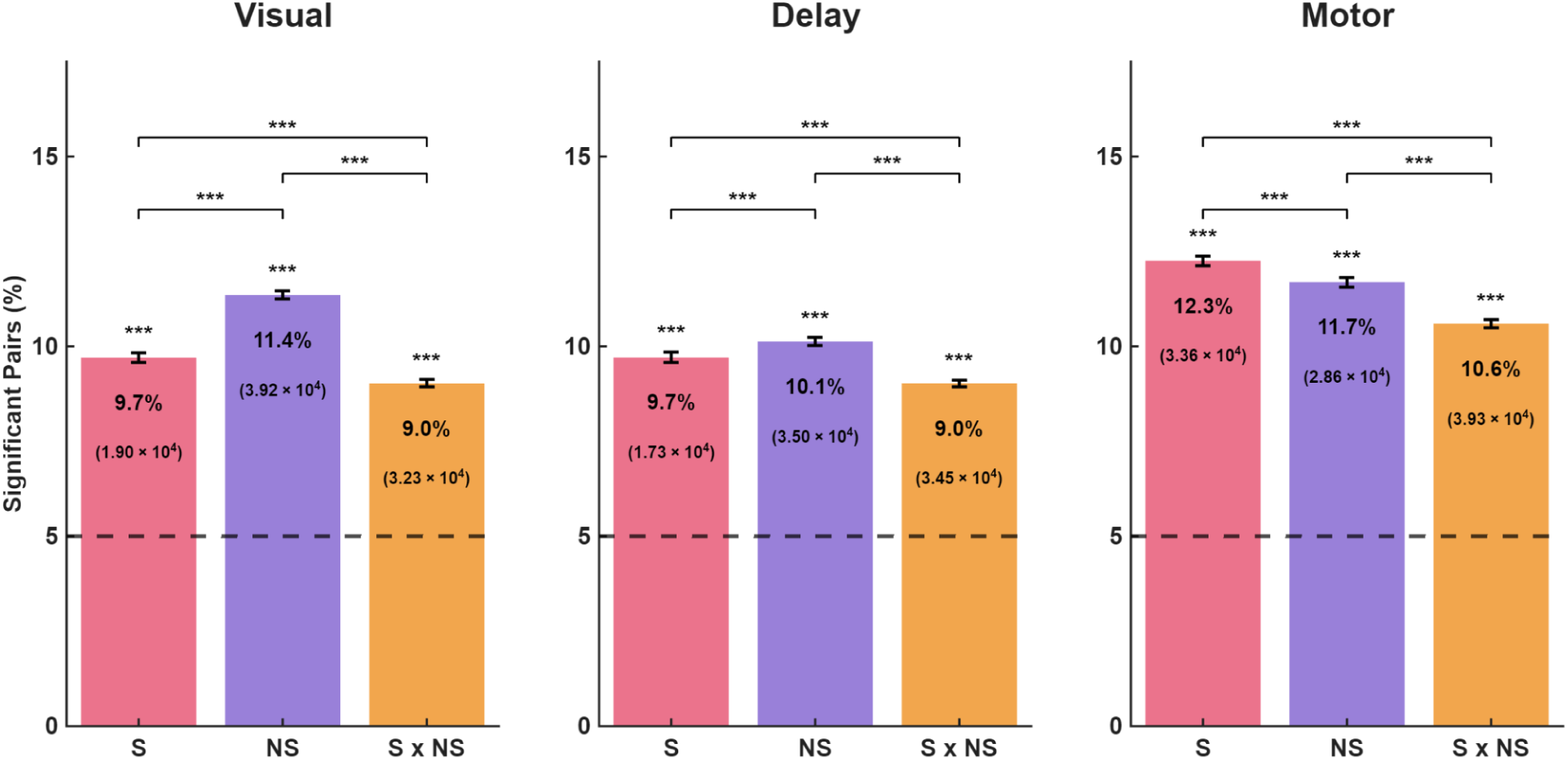
Condition-dependent NCs in selective, non-selective and mixed neuronal pairs. Bar graphs show the percentage of neuron pairs with condition-dependent NCs for selective (S), non-selective (NS), and mixed (S x NS) pairs. The percentage of significant pairs is shown within each bar, with the number of significant pairs indicated in parentheses. Data are shown for neuronal pairs within 1000 μm of one other and for activity measured during the Visual (left), Delay (middle), and Motor (right) epochs. Error bars denote the 95% binomial confidence intervals. The horizontal dashed line indicates 5% chance level, and asterisks directly above the error bars show significant deviations from this chance level (***, p < 0.001, Chi-square goodness-of-fit test). Brackets at the top indicate significant differences in the proportions between categories (***, p < 0.001, Chi-square test for equality of proportions).

## Discussion

In this study, we investigated the prevalence and spatial dependence of condition-dependent NCs in the LPFC during a spatial delayed response task composed of visual, memory, and motor epochs. Within simultaneous recordings from hundreds of neurons, we observed that a significant population of neuron pairs exhibited condition-dependent NCs across Visual, Delay, and Motor epochs. Critically, while previous studies have largely focused on condition-dependent NCs among neurons with SC selectivity, we observed condition-dependent NCs of comparable magnitude among nonselective neurons. These effects showed a strong spatial dependence: nearby pairs were more likely to exhibit condition-dependent NCs in both selective and nonselective neuron pairs. The results demonstrate that correlated variability in spiking activity can be condition-dependent even in the absence of condition-dependent SCs. Our findings suggest that NCs are not merely a byproduct of shared synaptic inputs among tuned neurons, but a robust feature of the population that can serve as a distinct channel for task-relevant information.

Our results describe condition-dependent NCs among local neuronal populations in the primate LPFC. The shared synaptic inputs presumably shaping these NCs are undetermined and may be local (Levitt et al., 1993; Kritzer and Goldman Rakic, 1995), subcortical (Fuster and Alexander, 1973; Funahashi et al., 2013; Guo et al., 2017), and/or intercortical in origin (Gnat & Andersen, 1988; Funahashi et al., 1989; Chafee & Goldman-Rakic, 1998; Froudist-Walsh et al., 2021). In recent years, high-density recordings have proven useful for identifying putative monosynaptic connections via short time-lag correlations in spike-timing (Trepka et al., 2022; Panichello et al., 2024; Zhu et al., 2025; Carr et al., 2026; Berenyi et al., 2013; Sibille et al., 2024 ; Sibille et al., 2022; Trautmann et al., 2025). Such an approach could prove useful for identifying shared inputs, for example, by examining whether neurons from a particular brain area disproportionately exhibit leading short time-lag correlations with LPFC neurons displaying condition-dependent NCs. These correlative approaches could be paired with causal manipulations (e.g., Merrikhi et al., 2017) to confirm monosynaptic connectivity. In addition to identifying common inputs shaping condition-dependent NCs, these tools can also be used to explore whether neurons with condition-dependent NCs have distinct patterns of efferent connectivity, suggesting a distinct computational role.

If condition-dependent NCs are indeed produced by tuned synaptic inputs, what explains the equal prevalence of conditioned-dependent NCs in nonselective neurons? One possibility is that these nonselective neurons may receive a large amount of inhibition and weaker tuned excitatory inputs. Intriguingly, application of a GABA antagonist to primate lateral prefrontal cortex during a spatial delayed response tasks elicits SC tuning in previously untuned neurons, consistent with inhibition masking spatially tuned excitatory inputs under normal conditions (Rao et al., 2000). Alternatively, inhibitory drive to nonselective cells may itself be tuned (Constantinidis & Goldman-Rakic, 2002; Rao et al., 1999). If neurons receive condition-independent excitatory inputs but condition-dependent inhibition, the resulting net fluctuations in neuronal excitability could similarly manifest as condition-dependent NCs without driving changes in SCs.

Additionally, it will be important to understand how much information about task-relevant variables the observed condition-dependent NCs can provide to downstream brain areas. From a population decoding perspective, this entails classifying task variables using the SC covariance across all simultaneously recorded neurons. The number of trials required for accurate covariance estimation, however, increases with the total number of neurons (Ledoit & Wolf, 2004). Accordingly, the number of training trials needed to keep test error within reasonable bounds for covariance-sensitive classification methods grows roughly quadratically with neuronal population size (Raudys & Jain, 1991). Obtaining such large trial counts will become increasingly tractable as methods for chronic high-density recordings (e.g., Angotzi et al., 2025) continue to improve, allowing experimental quantification of the amount of information provided by populations of NCs.

## Methods

### Animal Subjects

Data were collected from three male rhesus monkeys (Macaca mulatta), designated as A (11 years, 11 kg), H (12 years, 14 kg), and J (8 years, 12 kg), as previously reported (Panichello et al., 2024). All experimental protocols and surgical procedures were performed in accordance with the National Institutes of Health guidelines and received approval from the Stanford University Institutional Animal Care and Use Committee (IACUC).

### Experimental Setup and Behavioral Task

Visual stimuli were displayed on a VIEWPixx3D monitor (60 cm viewing distance) by MATLAB (R2022a) and Psychtoolbox. Eye movements were recorded at 1 kHz using an Eyelink 1000 system. Trials began with the monkeys fixating on a central spot against a grey background for 600-800 ms. Subsequently, a square cue (green, black, or white for monkey A, H, and J, respectively) measuring 1 degree of visual angle (DVA) on a side appeared for 50 ms at one of eight possible locations (5-7° eccentricity, 45° spacing). After a variable memory delay (1,400-1,600 ms), the fixation spot was extinguished, cuing the response phase, which took one of two forms. On Match-to-Sample (MTS) trials, two targets (blue circles, 1 DVA radius) were presented: one at the cued location and another at a distractor location (one out of the remaining seven locations). On Memory-Guided Saccade (MGS) trials, no targets were shown. Monkeys A and H were trained on and performed randomly interleaved MTS and MGS trials, whereas Monkey J was trained on and completed only MGS trials. To receive a fluid reward, subjects had to saccade to within 5 DVA of the cued location and maintain gaze for 200 ms. Strict fixation (within 2 DVA of the fixation mark for monkeys H and J, within 3 DVA for monkey A) was required throughout the trial up until the response. Intertrial intervals (ITI) were 300-1,000 ms after correct trials, while fixation breaks and incorrect responses resulted in a 2,000 ms ITI without reward.

### Surgical Procedures and Electrophysiology

Subjects were implanted with titanium headposts for head stabilization and recording chambers for cortical access. Recordings were obtained from area 8 and the principal sulcus (9/46). Specifically, recordings were obtained from area 8 in monkey A, areas 8 and 9/46 in monkey H, and area 9/46 in monkey J. Neural activity was recorded using primate Neuropixels probes (384 active channels, 3.84 mm span). Probes were lowered through the dura via a 21-gauge cannula using custom 3D-printed grids and motorized drives (NAN Instruments). To ensure stable recordings and mitigate probe drift, we allowed at least 30 minutes for the neurons to settle prior to data collection. Neural activity was monitored and saved using SpikeGLX (https://billkarsh.github.io/SpikeGLX/).

### Data Preprocessing and Analysis

Spike sorting was performed offline using Kilosort3, and both putative single- and multi-unit clusters were included in the analysis. Spike times were aligned to cue onset for each trial and binned with a width and timestep of 1 ms. Neurons that fired fewer than 1,000 spikes in the experimental sessions (each ∼3 hours) were excluded from analysis.

### Statistical Standards

Unless noted otherwise, all statistical tests were two-sided. Task locations were randomized within each session to ensure balanced exposure for each subject within each session. Neurons were recorded without selection bias, with electrode placement optimizing for signal-to-noise ratio.

### Neuronal Classification via Population Decoding

We used multi-class linear decoders (support vector machines, as implemented by the fitcecoc function in MATLAB) to assess the amount of information about the cue location in neuronal spike counts. This analysis was performed separately for three task epochs: Visual (0-300 ms post-cue onset), Memory (400-1,200 ms post-cue onset), and Motor (-125-0 ms relative to movement onset). For each session, spike counts were summed across each epoch on each trial and z-scored across all trials per neuron. Then, for each session and epoch, a decoder was trained to predict the cue location using a one-vs-one approach based on the vector of spike counts across the population. Overall classification performance was assessed using 10-fold cross-validation.

### Neuron Importance and Recursive Feature Ablation

To identify the contribution of individual neurons to the population representation, we analyzed the decoder weights. For each fold, we extracted the absolute weights (|W|) from all learners and averaged them across learners and folds to obtain an importance score for each neuron. We then performed a neuron dropping analysis (recursive feature ablation). Units were ranked by their importance scores and sequentially removed from the population. After each removal, the 10-fold cross-validated decoding accuracy was recalculated using the remaining subset of neurons. This iterative process continued until all neurons were removed, allowing us to trace the decay of decoding performance as a function of unit loss.To define a threshold for selectivity, we first calculated the 95% binomial confidence interval (CI) for the decoding accuracy at each removal step.

Next, we identified the point in the dropping analysis when decoding performance first dropped to chance level (i.e., when the CI included chance performance: 12.5% for 8 possible cue locations). All neurons removed prior to this point were classified as Selective, as their presence was necessary to maintain decoding accuracy above chance. Conversely, neurons remaining after this threshold were classified as Non-Selective, as their removal did not further degrade performance below the chance-level baseline. Some sessions never reached chance level and were therefore, not included in subsequent analysis. Finally, units were sorted into functional subclasses (e.g., Visual-selective, Delay-selective, or Motor selective) based on the specific epochs during which they met the selectivity criteria.

### Calculation of Pairwise Noise Correlations

To quantify noise correlations (r_sc_), spike counts for each unit and epoch were converted into z-scores by subtracting the mean and dividing by the standard deviation of spike counts across all correct trials for each cue location:

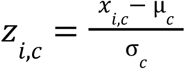

where *x_i,c_* is the spike count on trial i for cue c, and µ_c_ and σ_c_ are the mean and standard deviation of spike counts for that cue, respectively. The noise correlation for a neuron pair was then defined as the Pearson correlation coefficient (r) between their z-scored spike counts across all trials for a specific cue location and epoch. Units that never fired during the task epoch of interest were excluded from analysis.

To investigate the spatial structure of noise correlations, we computed the center of mass of the spike waveform template for each unit using Neuropixels Utils (https://djoshea.github.io/neuropixel-utils/) and calculated the Euclidean distance between neurons in each pair.

### Pairwise Linear Regression Analysis

To evaluate the cue-dependent modulation of noise correlations, we employed linear regression, using MATLAB’s fitlm function. For each simultaneously recorded neuron pair, we modeled the z-scored spike count of one neuron (N_2_) as a function of the spike count of the other (N_1_) and its interaction with the cue location (C). We used the model below:

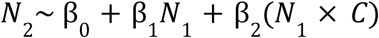

The model tested whether the slope of the relationship between N_1_ and N_2_ varied across the eight cue locations. Statistical significance of the interaction term was determined using an ANOVA (F-test) comparing the two models. All spike counts were z-scored within each session to account for differences in baseline excitability.

To quantify the prevalence of location-dependent modulation, the analysis was restricted to neuron pairs separated by less than 1000 μm. For these pairs, the number of neuron pairs showing a significant effect in the condition-dependent interaction term (p < 0.05) was divided by the total number of pairs for each neuron pair type (selective (S), non-selective (NS), and mixed (S x NS) pairs), and expressed as a percentage.

## Acknowledgments

We thank Danielle Lopes and Stephen Cital for technical assistance.

## References

Aliramezani, M., Constantinidis, C., & Daliri, M. R. (2025). Unraveling the roles of spatial working memory sustained and selective neurons in prefrontal cortex. Communications biology, 8(1), 767. 10.1038/s42003-025-08211-8

Angotzi, G. N., Baker, A. M. E., Vincenzi, M., Orban, G., Ribeiro, J. F., Tenorio, V., Berdondini, L., & Baker, S. N. (2025). High-density, Identified Cell Recordings from Motor Cortex of Awake Behaving Macaques using 1024-channel SiNAPS-NHP Probes. bioRxiv : the preprint server for biology, 2025.07.22.665434. 10.1101/2025.07.22.665434

Berényi, A., Somogyvári, Z., Nagy, A. J., Roux, L., Long, J. D., Fujisawa, S., Stark, E., Leonardo, A., Harris, T. D., & Buzsáki, G. (2014). Large-scale, high-density (up to 512 channels) recording of local circuits in behaving animals. Journal of neurophysiology, 111(5), 1132–1149. 10.1152/jn.00785.2013

Bondy, A. G., Haefner, R. M., & Cumming, B. G. (2018). Feedback determines the structure of correlated variability in primary visual cortex. Nature neuroscience, 21(4), 598–606. 10.1038/s41593-018-0089-1

Bruce, C. J., & Goldberg, M. E. (1985). Primate frontal eye fields. I. Single neurons discharging before saccades. Journal of neurophysiology, 53(3), 603–635. 10.1152/jn.1985.53.3.603

Carr, N., Zhu, S., Chen, X., Lee, E. K., Perliss, A., Moore, T., & Chandrasekaran, C. (2026). Neuropixels reveal laminar microcircuit organization in monkey V1 in vivo. Proceedings of the National Academy of Sciences of the United States of America, 123(8), e2521556123. 10.1073/pnas.2521556123

Chafee, M. V., & Goldman-Rakic, P. S. (1998). Matching patterns of activity in primate prefrontal area 8a and parietal area 7ip neurons during a spatial working memory task. Journal of neurophysiology, 79(6), 2919–2940. 10.1152/jn.1998.79.6.2919

Cohen, M. R., & Kohn, A. (2011). Measuring and interpreting neuronal correlations. Nature neuroscience, 14(7), 811–819. 10.1038/nn.2842

Downer, J. D., Rapone, B., Verhein, J., O’Connor, K. N., & Sutter, M. L. (2017). Feature-Selective Attention Adaptively Shifts Noise Correlations in Primary Auditory Cortex. The Journal of neuroscience : the official journal of the Society for Neuroscience, 37(21), 5378–5392. 10.1523/JNEUROSCI.3169-16.2017

Franke, F., Fiscella, M., Sevelev, M., Roska, B., Hierlemann, A., & da Silveira, R. A. (2016). Structures of Neural Correlation and How They Favor Coding. Neuron, 89(2), 409–422. 10.1016/j.neuron.2015.12.037

Froudist-Walsh, S., Bliss, D. P., Ding, X., Rapan, L., Niu, M., Knoblauch, K., Zilles, K., Kennedy, H., Palomero-Gallagher, N., & Wang, X. J. (2021). A dopamine gradient controls access to distributed working memory in the large-scale monkey cortex. Neuron, 109(21), 3500–3520.e13. 10.1016/j.neuron.2021.08.024

Funahashi S. (2013). Thalamic mediodorsal nucleus and its participation in spatial working memory processes: comparison with the prefrontal cortex. Frontiers in systems neuroscience, 7, 36. 10.3389/fnsys.2013.00036

Funahashi, S., Bruce, C. J., & Goldman-Rakic, P. S. (1989). Mnemonic coding of visual space in the monkey’s dorsolateral prefrontal cortex. Journal of neurophysiology, 61(2), 331–349. 10.1152/jn.1989.61.2.331

Fuster, J. M., & Alexander, G. E. (1973). Firing changes in cells of the nucleus medialis dorsalis associated with delayed response behavior. Brain research, 61, 79–91. 10.1016/0006-8993(73)90517-9

Gnadt, J. W., & Andersen, R. A. (1988). Memory related motor planning activity in posterior parietal cortex of macaque. Experimental brain research, 70(1), 216–220. 10.1007/BF00271862

Guo, Z. V., Inagaki, H. K., Daie, K., Druckmann, S., Gerfen, C. R., & Svoboda, K. (2017). Maintenance of persistent activity in a frontal thalamocortical loop. Nature, 545(7653), 181–186. 10.1038/nature22324

Hofer, S. B., Ko, H., Pichler, B., Vogelstein, J., Ros, H., Zeng, H., Lein, E., Lesica, N. A., & Mrsic-Flogel, T. D. (2011). Differential connectivity and response dynamics of excitatory and inhibitory neurons in visual cortex. Nature neuroscience, 14(8), 1045–1052. 10.1038/nn.2876

Hu, Y., Trousdale, J., Josić, K., & Shea-Brown, E. (2012). Motif statistics and spike correlations in neuronal networks. BMC Neuroscience, 13(Suppl 1), P43. 10.1186/1471-2202-13-S1-P43

Josić, K., Shea-Brown, E., Doiron, B., & de la Rocha, J. (2009). Stimulus-dependent correlations and population codes. Neural computation, 21(10), 2774–2804. 10.1162/neco.2009.10-08-879

Kerlin, A. M., Andermann, M. L., Berezovskii, V. K., & Reid, R. C. (2010). Broadly tuned response properties of diverse inhibitory neuron subtypes in mouse visual cortex. Neuron, 67(5), 858–871. 10.1016/j.neuron.2010.08.002

Kim, D., Jeong, H., Lee, J., Ghim, J. W., Her, E. S., Lee, S. H., & Jung, M. W. (2016). Distinct Roles of Parvalbumin- and Somatostatin-Expressing Interneurons in Working Memory. Neuron, 92(4), 902–915. 10.1016/j.neuron.2016.09.023

Kohn, A., & Smith, M. A. (2005). Stimulus dependence of neuronal correlation in primary visual cortex of the macaque. The Journal of neuroscience : the official journal of the Society for Neuroscience, 25(14), 3661–3673. 10.1523/JNEUROSCI.5106-04.2005

Kritzer, M. F., & Goldman-Rakic, P. S. (1995). Intrinsic circuit organization of the major layers and sublayers of the dorsolateral prefrontal cortex in the rhesus monkey. The Journal of comparative neurology, 359(1), 131–143. 10.1002/cne.903590109

Ledoit, O., & Wolf, M. (2004). A well-conditioned estimator for large-dimensional covariance matrices. Journal of multivariate analysis, 88(2), 365–411. 10.1016/S0047-259X(03)00096-4

Levitt, J. B., Lewis, D. A., Yoshioka, T., & Lund, J. S. (1993). Topography of pyramidal neuron intrinsic connections in macaque monkey prefrontal cortex (areas 9 and 46). The Journal of comparative neurology, 338(3), 360–376. 10.1002/cne.903380304

Merrikhi, Y., Clark, K., Albarran, E., Parsa, M., Zirnsak, M., Moore, T., & Noudoost, B. (2017). Spatial working memory alters the efficacy of input to visual cortex. Nature communications, 8, 15041. 10.1038/ncomms15041

Nassar, M. R., Scott, D., & Bhandari, A. (2021). Noise Correlations for Faster and More Robust Learning. The Journal of neuroscience : the official journal of the Society for Neuroscience, 41(31), 6740–6752. 10.1523/JNEUROSCI.3045-20.2021

Panzeri, S., Macke, J. H., Gross, J., & Kayser, C. (2015). Neural population coding: combining insights from microscopic and mass signals. Trends in cognitive sciences, 19(3), 162–172. 10.1016/j.tics.2015.01.002

Panzeri, S., Schultz, S. R., Treves, A., & Rolls, E. T. (1999). Correlations and the encoding of information in the nervous system. Proceedings. Biological sciences, 266(1423), 1001–1012. 10.1098/rspb.1999.0736

Pola, G., Thiele, A., Hoffmann, K. P., & Panzeri, S. (2003). An exact method to quantify the information transmitted by different mechanisms of correlational coding. Network (Bristol, England), 14(1), 35–60. 10.1088/0954-898x/14/1/303

Ponce-Alvarez, A., Thiele, A., Albright, T. D., Stoner, G. R., & Deco, G. (2013). Stimulus-dependent variability and noise correlations in cortical MT neurons. Proceedings of the National Academy of Sciences of the United States of America, 110(32), 13162–13167. 10.1073/pnas.1300098110

Qi, X. L., & Constantinidis, C. (2012). Correlated discharges in the primate prefrontal cortex before and after working memory training. The European journal of neuroscience, 36(11), 3538–3548. 10.1111/j.1460-9568.2012.08267.x

Rao, S. G., Williams, G. V., & Goldman-Rakic, P. S. (1999). Isodirectional tuning of adjacent interneurons and pyramidal cells during working memory: evidence for microcolumnar organization in PFC. Journal of neurophysiology, 81(4), 1903–1916. 10.1152/jn.1999.81.4.1903

Rao, S. G., Williams, G. V., & Goldman-Rakic, P. S. (2000). Destruction and creation of spatial tuning by disinhibition: GABA(A) blockade of prefrontal cortical neurons engaged by working memory. The Journal of neuroscience : the official journal of the Society for Neuroscience, 20(1), 485–494. 10.1523/JNEUROSCI.20-01-00485.2000

Raudys, S. J., & Jain, A. K. (1991). Small sample size effects in statistical pattern recognition: recommendations for practitioners. IEEE transactions on pattern analysis and machine intelligence, 13(3), 252–264. 10.1109/34.75512

Runyan, C. A., & Sur, M. (2013). Response selectivity is correlated to dendritic structure in parvalbumin-expressing inhibitory neurons in visual cortex. The Journal of neuroscience : the official journal of the Society for Neuroscience, 33(28), 11724–11733. 10.1523/JNEUROSCI.2196-12.2013

Scaglione, A., Moxon, K. A., Aguilar, J., & Foffani, G. (2011). Trial-to-trial variability in the responses of neurons carries information about stimulus location in the rat whisker thalamus. Proceedings of the National Academy of Sciences of the United States of America, 108(36), 14956–14961. 10.1073/pnas.1103168108

Scholl, B., Pattadkal, J. J., Dilly, G. A., Priebe, N. J., & Zemelman, B. V. (2015). Local Integration Accounts for Weak Selectivity of Mouse Neocortical Parvalbumin Interneurons. Neuron, 87(2), 424–436. 10.1016/j.neuron.2015.06.030

Shamir, M., & Sompolinsky, H. (2004). Nonlinear population codes. Neural computation, 16(6), 1105–1136. 10.1162/089976604773717559

Shapcott, K. A., Schmiedt, J. T., Saunders, R. C., Maier, A., Leopold, D. A., & Schmid, M. C. (2016). Correlated activity of cortical neurons survives extensive removal of feedforward sensory input. Scientific reports, 6, 34886. 10.1038/srep34886

Shilling-Scrivo, K., Mittelstadt, J., & Kanold, P. O. (2022). Decreased Modulation of Population Correlations in Auditory Cortex Is Associated with Decreased Auditory Detection Performance in Old Mice. The Journal of neuroscience : the official journal of the Society for Neuroscience, 42(49), 9278–9292. 10.1523/JNEUROSCI.0955-22.2022

Sibille, J., Gehr, C., Benichov, J. I., Balasubramanian, H., Teh, K. L., Lupashina, T., Vallentin, D., & Kremkow, J. (2022). High-density electrode recordings reveal strong and specific connections between retinal ganglion cells and midbrain neurons. Nature communications, 13(1), 5218. 10.1038/s41467-022-32775-2

Sibille, J., Gehr, C., & Kremkow, J. (2024). Efficient mapping of the thalamocortical monosynaptic connectivity in vivo by tangential insertions of high-density electrodes in the cortex. Proceedings of the National Academy of Sciences of the United States of America, 121(4), e2313048121. 10.1073/pnas.2313048121

Singer W. (1999). Neuronal synchrony: a versatile code for the definition of relations?. Neuron, 24(1), 49–125. 10.1016/s0896-6273(00)80821-1

Smith, M. A., & Kohn, A. (2008). Spatial and temporal scales of neuronal correlation in primary visual cortex. The Journal of neuroscience : the official journal of the Society for Neuroscience, 28(48), 12591–12603. 10.1523/JNEUROSCI.2929-08.2008

Trautmann EM, Hesse JK, Stine GM, Xia R, Zhu S, O’Shea DJ, Karsh B, Colonell J, Lanfranchi FF, Vyas S, Zimnik A, Amematsro E, Steinemann NA, Wagenaar DA, Pachitariu M, Andrei A, Lopez CM, O’Callaghan J, Putzeys J, Raducanu BC, Welkenhuysen M, Churchland M, Moore T, Shadlen M, Shenoy K, Tsao D, Dutta B, Harris T. (2025). Large-scale high-density brain-wide neural recording in nonhuman primates. Nat Neurosci, *28*(7):1562-1575. 10.1038/s41593-025-01976-5

Trepka, E. B., Zhu, S., Xia, R., Chen, X., & Moore, T. (2022). Functional interactions among neurons within single columns of macaque V1. eLife, 11, e79322. 10.7554/eLife.79322

Vaadia, E., Haalman, I., Abeles, M., Bergman, H., Prut, Y., Slovin, H., & Aertsen, A. (1995). Dynamics of neuronal interactions in monkey cortex in relation to behavioural events. Nature, 373(6514), 515–518. 10.1038/373515a0

Wood, K. C., Blackwell, J. M., & Geffen, M. N. (2018). Cortical inhibitory interneurons control sensory processing. Current opinion in neurobiology, 46, 200–207. 10.1016/j.conb.2017.08.018

Zavitz, E., & Price, N. S. C. (2019). Understanding Sensory Information Processing Through Simultaneous Multi-area Population Recordings. Frontiers in neural circuits, 12, 115. 10.3389/fncir.2018.00115

Zhu, S., Lopes, D. A., Cital, S. N., & Moore, T. (2026). Ensemble Coding of Hidden Objects in Visual Cortex. bioRxiv : the preprint server for biology, 2025.12.29.696959. 10.64898/2025.12.29.696959

Zylberberg, J., Cafaro, J., Turner, M. H., Shea-Brown, E., & Rieke, F. (2016). Direction-Selective Circuits Shape Noise to Ensure a Precise Population Code. Neuron, 89(2), 369–383. 10.1016/j.neuron.2015.11.019

